# Multimodal sensorimotor integration of visual and kinaesthetic afferents modulates motor circuits in humans

**DOI:** 10.1101/2021.01.31.429002

**Authors:** Volker R. Zschorlich, Frank Behrendt, Marc H.E. de Lussanet

## Abstract

Optimal motor control requires the effective integration of multi-modal information. Visual information of movement performed by others even enhances potentials in the upper motor neurons, through the mirror-neuron system. On the other hand, it is known that motor control is intimately associated with afferent proprioceptive information. Kinaesthetic information is also generated by passive, external-driven movements. In the context of sensory integration, its an important question, how such passive kinaesthetic information and visually perceived movements are integrated. We studied the effects of visual and kinaesthetic information in combination, as well as isolated, on sensorimotor-integration – compared to a control condition. For this, we measured the change in the excitability of motor cortex (M1) using low-intensity TMS. We hypothesised that both visual motoneurons and kinaesthetic motoneurons could enhance the excitability of motor responses. We found that passive wrist movements increase the motor excitability, suggesting that kinaesthetic motoneurons do exist. The kinaesthetic influence on the motor threshold was even stronger than the visual information. Moreover, the simultaneous visual and passive kinaesthetic information increased the cortical excitability more than each of them independently. Thus, for the first time, we found evidence for the integration of passive kinaesthetic- and visual-sensory stimuli.

## 1. Introduction

Sensorimotor control processes of targeted movements in sport and music requires effective performance in the integration of different sensory modalities. One aspect of these operations is summarized as multi-sensory integration and has been addressed in various paradigms.

With regard to multimodal information processing, it has been assumed that visual information usually plays a dominant role [1], [2], but the visual modality does not always dominate the other senses during cross-modal information processing [3]. Ernst and Bülthoff [4] found that the merging of information is following the principle of maximum likelihood [5], [6]. The brain combines information according to least variance and greatest reliability and weighs the information of corresponding sensors to achieve a robust percept and motor control.

One method to investigate the interaction between visual sensory information and motor circuits on a neuronal level is the measure of single neuron activity. This was carried out in the premotor cortices in non-human primates [7] and, more recently, even in the human brain [8]. Nerve cells that associate visual information in the context of movements with the activation of the same motor actions are referred to as mirror neurons. Typical electrophysiological recordings of such mirror neurons in monkeys [7], [9], [10], [11] have shown a discharge both during execution and perception of actions. Since these integration processes involve numerous neurons in the motor cortex, the influence of visual information on motor circuits can also be studied using transcranial magnetic stimulation (TMS) [12], [13], [14], [15]. The visual perception of motion activates sensory- and motor-representations in the brain [16], [17], [18], [19]. These brain regions are thought to play a central role in movement perception and movement generation as there is an activation during action execution and during action recognition [9], [20], [21].

We propose that an analogous functional property exists for *kinaesthetic* neurons. Kinaesthetic information provides the brain with a sense of movement and position of the limbs in relation to the whole body. Kinaesthetic information comes from proprioceptors, like muscle spindles and Golgi-tendon organs, and other receptors that we are largely unaware of (for review see [22]). With the arrival of coherent information of different modalities, motor circuits respond to an TMS with an elevated motor evoked potential (MEP). The processing of such multimodal information leads to a more precise and differentiated routing of the circuits in motor preparation processes. This, in turn, leads to a more pronounced corticospinal excitability [23]. It has already been shown that visually presented actions can cause such an increase in cortico-spinal excitability [14], [15]. On the other hand, on the basis of the hypothesis of dedifferentiation in old age [24], [25], [26] it was predicted and found that the process of motor preparation should lead to less pronounced MEP responses from motor cortex stimulation in elderly subjects [27]. For example, in young, non-human primates, it was only possible to initiate a targeted triggering of MEPs after the development of advanced motor skills, which are accompanied by a differentiation of cortical motor structures in ontogenesis [28], [29].

An important question is, how the different sensory modalities are related, and how they are brought into a common “data format”. Putative multi-modality neurons [30], [31], [32], [33] in motor-related cortices could take over this task of multisensory-integration projecting to the primary motor cortex. More recently, a significant role in sensorimotor integration has been attributed to the striatum [34]. If kinaesthetic- and visual information are processed in the same way during motor preparation, we proposed above, both, seeing a movement and feeling a passive movement should enhance the corticospinal excitability.

In the present experiment kinaesthetic sensory information was provided by a motor-driven motion of the wrist. The visual information of wrist movement was given by a video clip of a moving hand executing a wrist extension or a wrist flexion movement. Through selective inclusion and exclusion of certain sensory modalities, the effects of visual and kinaesthetic information on motor preparation processes can be analysed. We hypothesize in this study, that additional sensory information elevates the MEP amplitudes so that inputs from two different modalities produces motor responses with higher response amplitudes as in the case of only a single modality or control condition.

## 2. Materials and Methods

### 2.1. Subjects

Twenty healthy participants with a mean age of 28.3 years (± 8.3) with no history of neurological disease or upper limb impairment were included. Participants had the opportunity to familiarize themselves with the experiment and with the treatment. In accordance with the Declaration of Helsinki ethical approval for the stimulation experiment was given from the ethics committee of the University Rostock – Germany. Volunteers gave their written consent to participate in the investigation.

### 2.2 Protocol

A total of 60 stimulations were applied for the three conditions: kinaesthetic in an extension movement (KIN), visual display of an extension movement (VIS), and as the third condition the combination of visual and kinaesthetic extension afferent information (VIS & KIN). For the control condition (CC) 20 Stimulation were applied. The magnetic stimulation was performed within an interval of minimal 5 s temporal extent. Volunteers were told to feel their wrist joint moving by looking at the monitor with the extending wrist joint in the VIS, KIN and in the VIS & KIN condition. All conditions were supplied in a random order. Coil placement and stimulation period lasted for a total of about 15 min. The CC measurement was conducted in a relaxed state without any movement looking on a small cross in the middle of the black screen instead of the displayed hand movement video clip (Figure 1).

**Figure 1.**
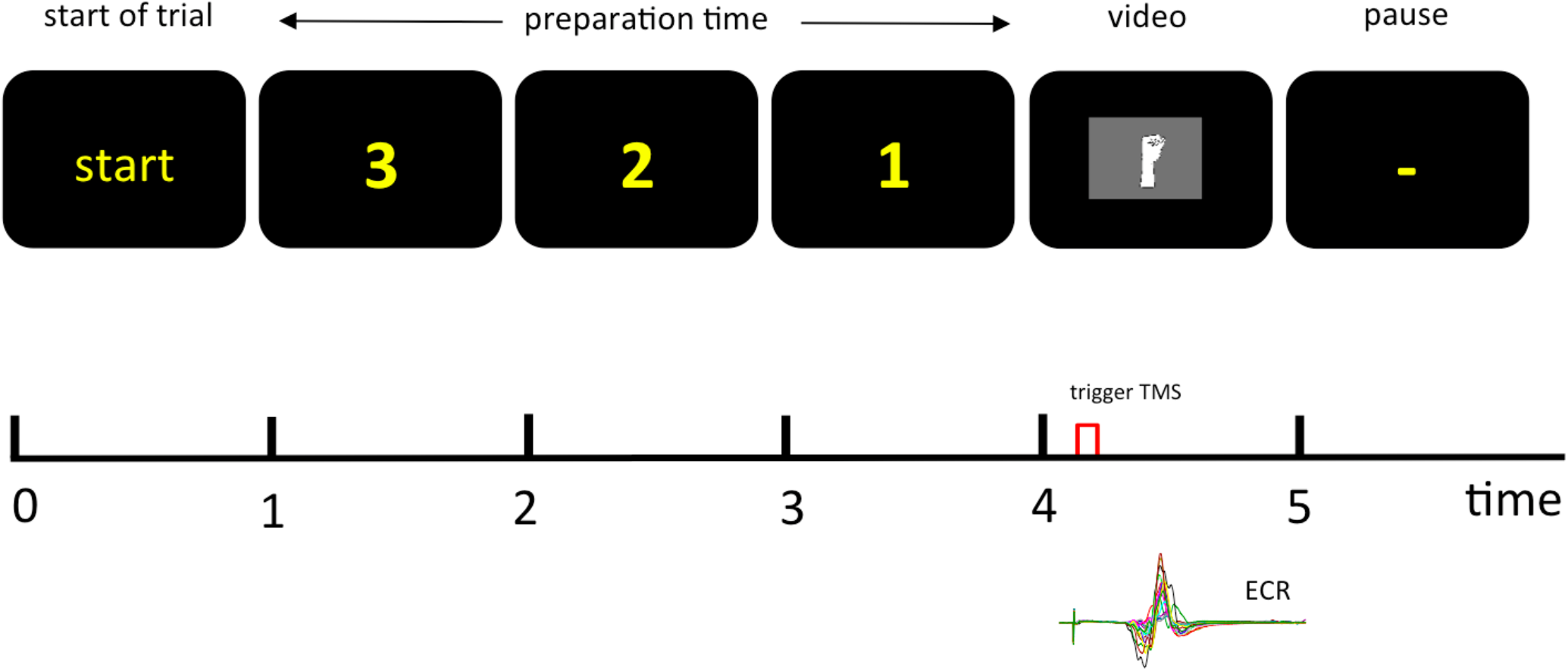
Schematic representation of the experimental set up shows the screen presentation for the different experimental conditions of sensory input. After the presentation of the start signal, there is a 3-second delay before the hand (VIS, VIS & KIN) was presented. In the kinaesthetic only condition (KIN), a black screen was shown instead. In the control condition (CC), a small fixation cross was shown. TMS was applied in the zero position during moving the wrist from the flexed to the extended position. The EMG of the extensor carpi radialis muscle (ECR) was measured.

### 2.3. Brain Stimulation

With TMS it is feasible to show the influence of different sensory input modes on the target-directed motor circuits of the human primary motor cortex [12], [13], [14], [15] and it is, therefore, possible to address some aspects of sensorimotor integration. Low-intensity transcranial stimuli [35], [36] were delivered slightly above motor threshold. With this method, it is possible to excite already subliminal elevated neurons in the primary motor cortex (M1) above threshold and to release an MEP. With low-intensity magnetic impulses it is viable to do a readout of the near threshold elevated upper motor neurons. For TMS a magnetic stimulator R30 MagPro with MagOption (MagVenture, Skovlunde Denmark - formerly Medtronic) and a parabolic coil type MMC-140 were used. A biphasic symmetric pulse of a duration of 280 μs was administered for stimulation. A biphasic pulse show more efficacy to activate neural tissue of the hand area [37], [38]. The coil was placed over the brain area where the lowest motor threshold could be found for the recorded muscle. Therefore, the stimulation side was about one to two cm behind the vertex over the right motor cortex with the concave side of the coil was placed to the surface of the skull. The coil was oriented over the right motor cortex. The coil position was kept constant with the help of two adjustable arms (Manfrotto Feltre, Italy) during the whole experiment. The magnetic gradient of the stimulator was defined as 20 *%* above the resting motor threshold to get sufficient detectable MEP on the left hand extensor carpi radialis muscle (ECR). The motor threshold was defined as stimulus intensity that produced an MEP of 100 μV in 3 of 5 trials. The used intensities just marginally above motor threshold were chosen for better detection the modulation effects of different senses on the motor preparatory processes. It could be shown that stimulation with 1.2 x resting motor threshold is suitable [39], [40]. The magnetic gradient in all trials lies therefore in the range between 45 – 75 A / μs.

### 2.4. Electromyography (EMG)

The TMS elicited motor evoked potentials were recorded with a custom made differential amplifier with an input resistance of 16 gΩ and a bandwidth of 1 Hz to 1000 Hz. The gain was chosen with a factor of 1000. Before calculation, EMG signals were high-pass filtered by a digital 1st order Butterworth filter with a cut-off frequency of 1 Hz [41]. The MEP responses were registered with two Ag-AgCl cup electrodes (Hellige baby-electrodes; GE medical systems, Milwaukee, USA) with an electrode surface area of *3 mm^2^* and were placed with a distance of 1 cm longitudinally over the belly of the extensor carpi radialis muscle. The skin preparation before electrode application was carried out with alcohol and the hairs were removed prior to skin abrasion [42].

The skin impedance at 30 Hz was always lower than Z = 10 kΩ. For conductance, an electrode gel (Parker Laboratories, Fairfield, USA) was used. Electrodes and twisted cables were fixed with self-adhesive tape on the skin. As reference electrode served an ECG-limb-clamp fixed at the upper arm located at the m. biceps. Muscle relaxation was strongly recommended and monitored visually over the whole experiment (Figure 2). MEPs were discarded offline, whenever background EMG amplitude higher than 50 μV was detectable in a time-window before 300 ms to movement onset. These events indicate a voluntary muscular activation, which has a prominent impact on MEP amplitudes [43]. The MEP were quantitatively evaluated as peak-to-peak amplitudes MEPpp [44], [45].

**Figure 2.**
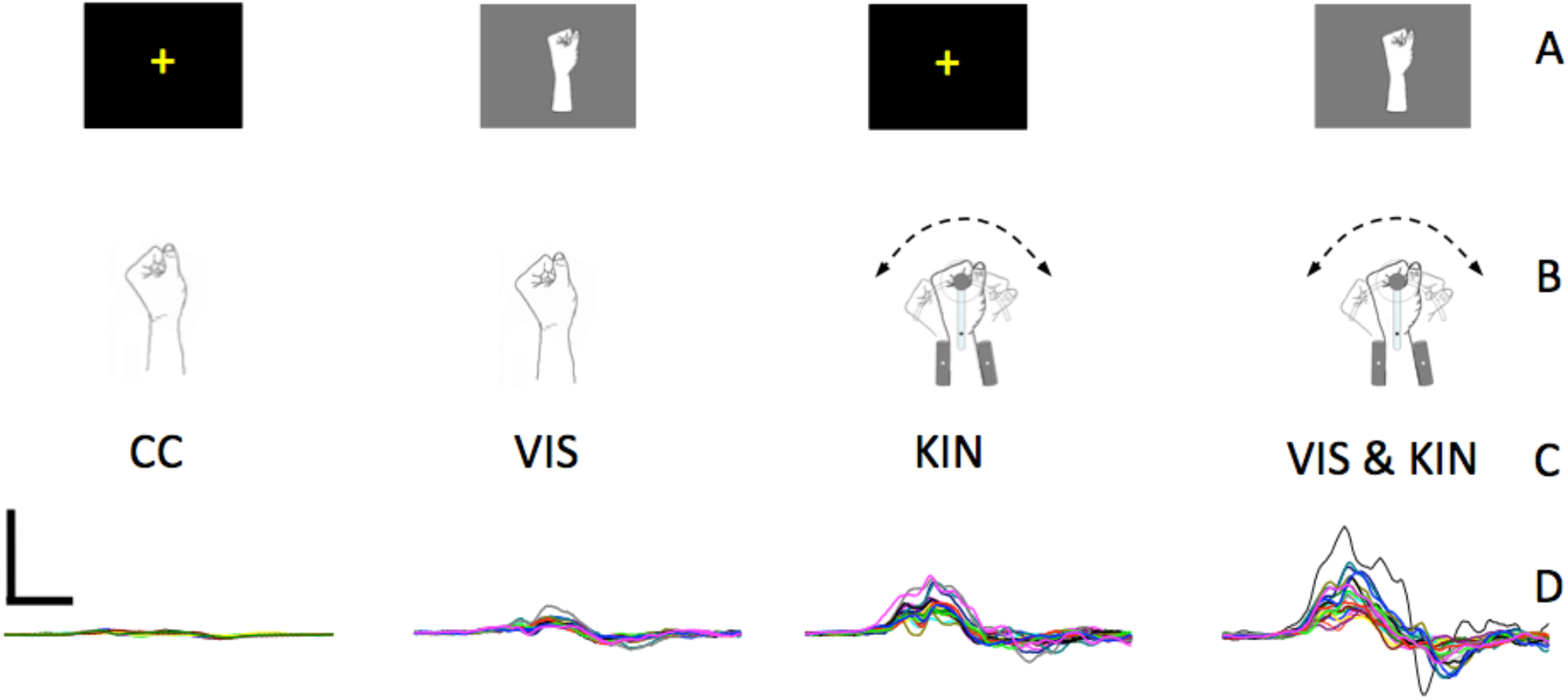
Representative trials of one subject illustrate the MEP of the extension directed movement. The different conditions represent the MEP response of the extensor carpi radialis at the control condition (CC), the visual wrist extension perception (VIS), the merely kinematic condition (KIN) and the coupled visual and kinematic wrist extension condition VIS & KIN (right). All MEPs have the same scale-units. Calibration bar: 0.5 mV, 10ms. A: Screen, B: Manipulandum, C: Condition, D: MEP.

### 2.5. Visual Stimuli

The hand in the presented animated drawing moves in the experimental KIN condition from the neutral position (0°) to an extension position (−30°) of the wrist joint with an angular velocity of 62,5°/s and subsequent duration of 480 ms. At this wrist extension position, the motor stops and a wait state of 200 ms followed. From this hand-extension position (−30°) the hand was moved to a flexion position of +30° with the same angular-velocity of 62,5°/s and a phase duration of 960 ms for the flexion whole movement (Figure 3). The movement executions carried out before the actual extension movement serves for embodiment or synchronization of the afferents. The following extension movement – relevant for the measurement – starting from this +30° position of the wrist and then moves with a speed of 62,5°/s into the extension.

**Figure 3.**
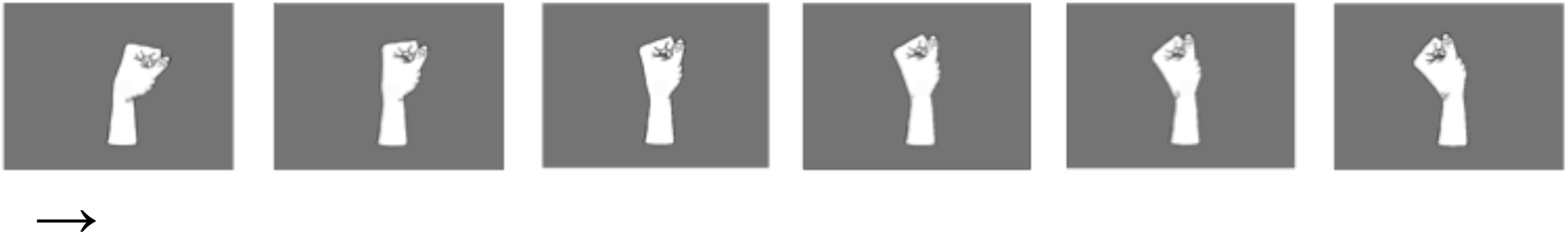
Screenshots of the hand in extension movement clip presented to the participants in the condition VIS- and VIS & KIN.

During this extension-movement, the stimulation was administered exactly at 0° to keep the corticospinal excitability independent from Ia spindle length [46]. The stimulation happens exactly 480 ms after starting the passive movement in the extension direction so that visual and kinaesthetic afferents can reach the motor cortex both. After a wait state of 200 ms in the extension position (−30°), the hand moves to the neutral position and stops there and the next randomized stimulation condition VIS, KIN or VIS & KIN can start. The visual demonstrated movement is exactly the same as the rotary actuator movement. The embodiment of the hand movement was achieved by giving the volunteers the opportunity of a sufficiently long preparation (minimal three minutes) for the stimulating conditions by visual presented as well as executed motions of the hand. In all condition, the participants were instructed to focus both on the feeling of the passive moving hand and on the moving hand on the video screen. For the visual stimulation in the VIS and in the VIS & KIN condition, the subjects were shown a video animation of an artificial hand motion on a monitor.

A stimulation at the view on the centre cross of the monitor was recorded in the CC and KIN condition. The control condition should help to exclude a possible modulation effect due to changes in attention. For this purpose, a small cross was presented in the middle of the monitor. The distance between the forehead-rest to the monitor was 110 cm and the display had an image size of 33*24 cm (about 18° angle of view).

### 2.6. Kinaesthetic stimuli

The kinaesthetic perception of the wrist motion was induced by passive driven limb movement. The subject grasped a handle with the left hand without any muscle activity and rests on a moveable support while the lower arm was fixed with a cast-construction (Figure 4). The centre alignment between extension and flexion was defined in a relaxed position where the extensor carpi radialis has no muscle activities. Subjects were sitting upright on a chair in front of the motor-driven wrist flexion-extension-machine. The left arm rested in a comfortable abduction of the upper arm and the forearm supported in a relaxed position between pronation and supination. The head of the volunteer was stabilized with a forehead-chin-rest to reduce head-to-coil movements. The left arm was bent with about 90 degree at the elbow joint, while the dominant (right) hand rested unrestrained on the table.

**Figure 4.**
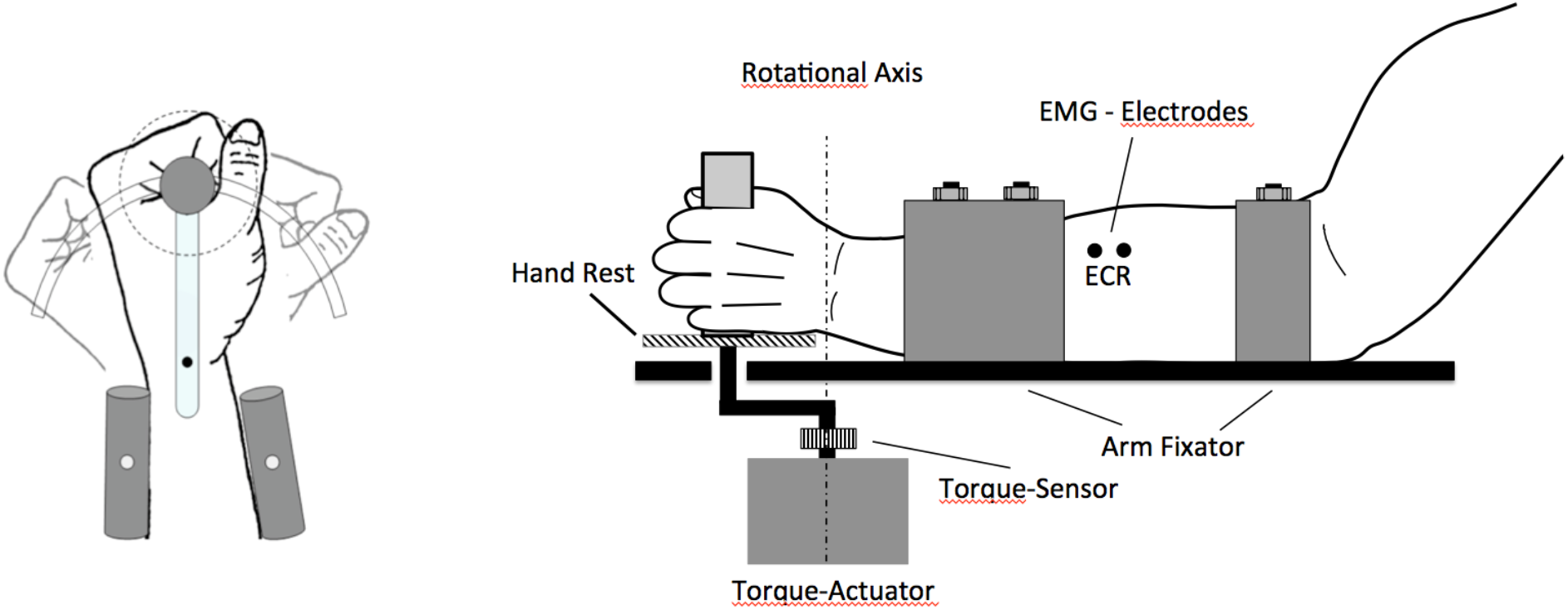
The graphical representation of the flexion-extension-machine for wrist movement shows a rotary motor driven lever, a hand rest (dotted circle on the right) and a forearm rest, and the moving hand grip. Electrodes were placed over the extensor carpi radialis (ECR).

The left-hand wrist joint movement was provided by a computer controlled brushless stepper motor (Copley Controls, Canton, MA 02021, U.S.A. type 145ST4M with 500 rpm). The rotary actuator (145ST4M, ALXION, France) was controlled by a Copley Xenus servo controller (type XTL-230-36/18-R) from the home position at neutral zero of the wrist. The moving of the crank arm covers an angle of 30 degrees (0.5235 rad) to maximum deflection. The speed of the crank arm had an angular velocity of 62.5 deg / s (1.09 rad/s) of the wrist joint extension or flexion. The given angular-velocity was achieved in less than 10 ms. The resolution of the resolver was 98260 steps for one revolution (2π rad).

To avoid injury, the range of motor movement was mechanically limited to ± 45° and a rapid shutdown was ensured for both the experimenter and the subjects. Care was taken to offer the stimuli in a synchronous manner otherwise the merging to a coherent percept could be disrupted or perturbed and a kind of de-afferents emerge [47]. The phase difference between the video clip and the torque actuator was less than one display frame throughout the complete movement sequence. In the crossmodal condition, the passive movement was coupled computer-controlled to a video sequence of the mentioned wrist movement. It was carefully controlled that the machine induced motion results only in an extension/flexion of the wrist and no movement has taken place on the forearm. The subject’s forearm and the actuator were hidden under a curtain. The visual demonstration was directly coupled to the actuator supported passive wrist movement with the same movement duration and angular velocity of the actuator supported movement.

### 2.7. Measurement and Data processing

All Data were sampled simultaneously with a 12 bit analog-digital-converter (DAQ-Card 6024 National Instruments, Austin, Texas, USA) at a sampling rate of 10000/s. DIAdem (National Instruments, Austin, Texas, USA) was used for data acquisition and further signal processing.

#### Statistics

Statistical analysis was performed using SPSS version 22.0 (SPSS Inc., Chicago, IL, USA) for windows. Peak-to-peak data (MEPpp) of the motor evoked potentials were calculated. This procedure was used to get a measure of corticospinal excitability. The absolute difference between the maximum and minimum value is expressed as the value of individual MEP.

An analysis of variance (ANOVA) for repeated measurement was calculated to test for significant differences of the experimental conditions. A post-hoc test was applied to test the significant difference between the demonstrated visual extension movement (VIS), the kinaesthetic extension movement (KIN), the third condition the synchronized combination of visual and kinaesthetic information (VIS & KIN), and the control condition (CC). The LSD correction was applied for multiple pairwise comparison. For correction, this test, in contrast to the simple paired comparison, the variances are calculated over all groups. The significance level was set to p < 0.05. The Greenhouse-Geisser coefficient (Ԑ) was employed to correct the degrees of freedom.

### 3. Results

In the current experiment, the ANOVA for repeated measurements shows a significant difference across the four conditions [*F*(1.25, 23.88) = 14.36_ε=0.42_; *p* < 0.000, η_p_^2^ = 0.43]. The lowest MEP values (0.49 mV ± 0.33mV) were measured in the control condition (CC), in which the test subjects only had to fix a small, stationary character in the middle of a black screen.

Depending on the offered modes of the movement-associated afferents, the MEPs were elevated. During observation of the wrist extension movement on the screen (VIS), the evoked MEP responses of the extensor carpi radialis muscle were larger as in the CC. With the VIS condition, we found the excitability of the MEPs by 0.65 mV ± 0.41 mV peak-to-peak values. In the kinaesthetic condition (KIN) the MEPs were elevated further and were larger than in the visual condition (VIS). The KIN condition showed an excitability of the MEPs by 1.13 mV ± 0.89 mV peak-to-peak values.

Coupled kinaesthetic information and visual information for the extension movement were measured in the forth condition (VIS & KIN). In the coupled condition VIS & KIN a further increase of the MEPs by 1.31 mV ± 0.89 mV peak-to-peak values appeared. The simultaneous observation of a wrist extension and kinaesthetic information of the passive extension in the VIS & KIN condition enhances the elicited MEPs compared to each solitary afferents and the CC (Figure 5).

**Figure 5.**
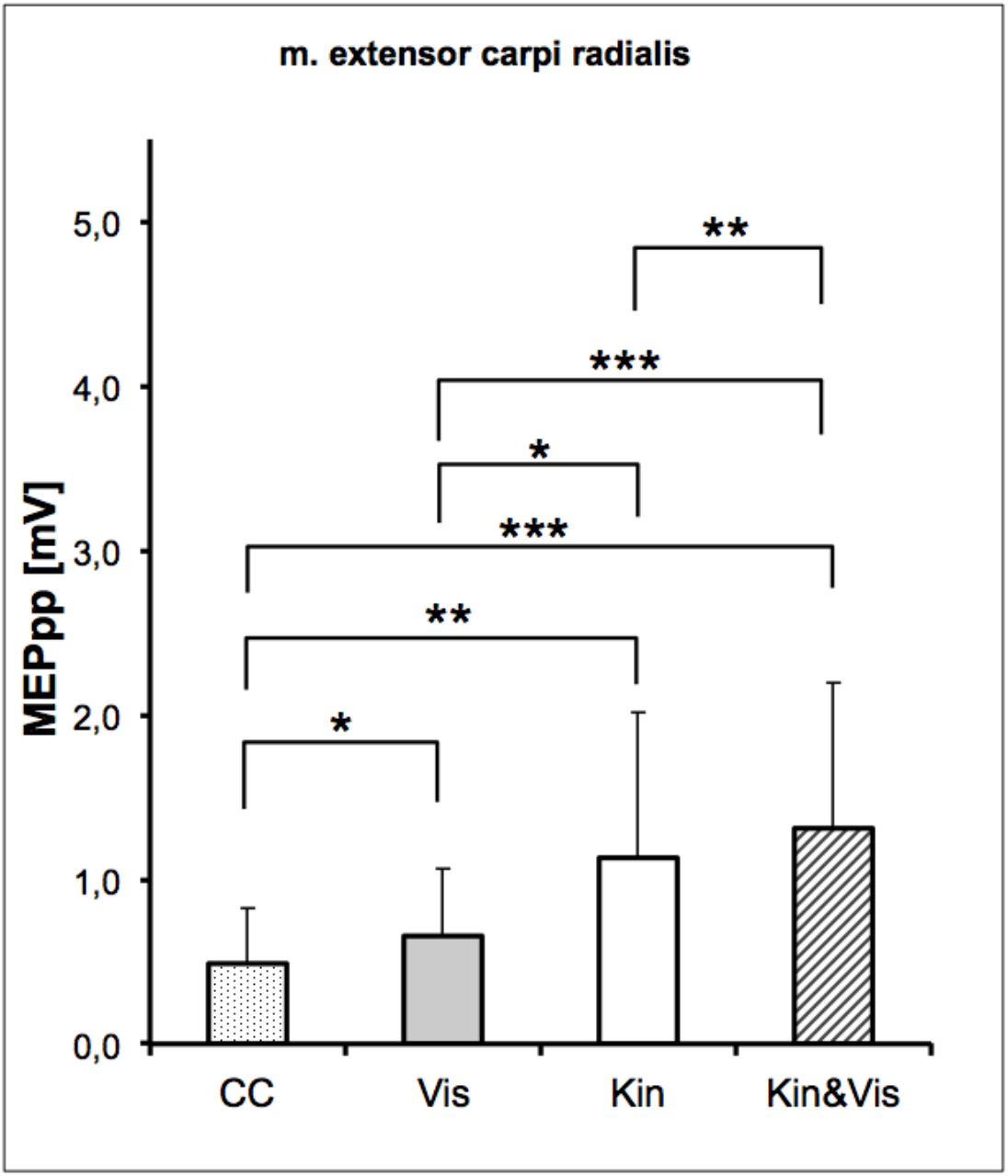
Mean MEP (± SD) across subjects, as a function of the four experimental conditions. Post-hoc comparisons of the MEP between all conditions shows significant differences. The MEP amplitudes shows an accumulation over the three different movement related modalities. The MEP amplitudes were presented as the mean ± standard deviation (SD). Significant differences in the MEP are presented with * *P* < 0.05, ** *P* < 0.01, and *** *P* < 0.001.

The LSD post-hoc test for the pairwise comparison of differences between the factors showed for all conditions a significant difference. The results of the paired comparison for each condition were presented in Table 1. The greatest difference was detected between the CC and the VIS & KIN condition. A dominant influence of kinaesthetic and visual motor information on motor preparation is evident. In all subjects, we found a modulation effect off the sensory integration on the corticospinal excitability.

**Table 1.** This table shows the post hoc results of the LSD pairwise comparisons between the 3 stimulation conditions and the control condition. We found for all comparisons significant differences. The Least Significant Difference Test is based on the two-sample t-test.

| Condition | Control | Visual | KIN |
| --- | --- | --- | --- |
| VIS | 0.015687<br>* |  |  |
| KIN | 0.001596<br>** | 0.015454<br>* |  |
| VIS & KIN | 0.000114<br>*** | 0.00081<br>*** | 0.004782<br>** |

## 4. Discussion

In the present TMS study, we explored the multimodal visual-proprioceptive interaction during the perception of a passive movement of the observer’s own hand. The data show that the evoked responses in the bimodal condition were significantly higher than in both unimodal conditions alone. The finding suggests that bimodal action perception has an additive influence, resulting in a more pronounced response to TMS when both modalities are presented simultaneously.

Nearly all events that we are exposed to every day presumably entail more than a single information modality and are thus of a multimodal character [48]. It is commonly accepted that a single human information-processing system alone is not capable of perceiving an event accurately enough in all circumstances and that an extensive percept can only be achieved by the integration of multiple sources of sensory information [4], [49]. Accordingly, several sources of information are permanently being combined, further processed and interpreted based on prior knowledge and experience. Classically, the visual perception was often considered the dominant one [50] even independent of others, but there have long been contradictory results [51]. The sense of movement and position of our body or limbs which is referred to as kinaesthesia [52] relies on proprioceptive afferents. This afferent information interacts with other senses such as vision in the perception process [53]. It was recently confirmed in a study on the contributions of visual and proprioceptive afferents to the perception of a passive arm displacement in virtual reality, that this process indeed involves non-visual signals and also that the latter interact with visual signals [54]. In an investigation of mental imagery on neuronal circuits of motor processing, it was found [55] that only kinaesthetic imagery, but not visual imagery, facilitates motor neurons in M1. This supports our results that kinaesthetic information is more effective than visual information in motor control processes. Moreover, Giroux et al. [54] reported that the interaction of the modalities had a strengthening effect on the vividness of the perception of the passive arm movement. This is in line with our results despite the fact that the authors did not test for the corticospinal excitability during the perception of the arm displacement but assessed speed and duration of the perceived movement verbally rated by the participants.

In a different experiment, also using TMS, it was found that there is an additive interaction of sound and sight during action observation [56]. Here, the authors used unimodal visual, unimodal auditory or both kinds of stimuli combined of hand action and also found a response increase in the multimodal condition. From the results it was concluded that there are likely no different kinds of action representations, i.e. a single representation for every modality, but rather shared action representations resulting in stronger responses if stimuli are perceived simultaneously and congruently. Our results support the hypothesis that action representations themselves comprise integrated multimodal information leading to an enhanced response to TMS in case action perception is multimodal and therefore more reliable.

The present study shows that kinaesthetic information affects motor circuits in M1. This means that motor nerve cells exist in M1, which connects kinaesthetic information with the associated excitatory motor structures. On a neuronal level the influence of visual movement information to the premotor cortex has been known already for over two decades [7]. It is therefore likely that significant portions of the processes of sensorimotor integration, which also pertains to cross-modal processing, are carried out putative in the premotor cortex (PMC) and M1 area. Strick and colleagues [57], [58] already proposed, that different input modalities in different locations of M1 represent task demands of high-level sensorimotor integration. This integration process may lead to a coherent percept. From studies of non-human primates [30], [31], [32] it was supposed that sensorimotor-integration is caused by multimodal neurons. Trimodal neurons in the ventral premotor cortex (PMv) were proposed [30] and Makin et al. [33] proposed involved trimodal neurons in PMC and intra-parietal sulcus (IPS) where visual, tactile and proprioceptive afferents are integrated within one neuron. More recently experiments found further brain areas, such as the lateral prefrontal cortex (lPFC) and the intraparietal sulcus (IPS), that provides resources for specialized motor control in dance and music [59]. An additional spinal contribution to the modulation of the MEPs cannot be completely ruled out, because a modulated discharge rate of spindle afferents may be crucial when the muscle is progressively lengthened [60], [61], [62]. However, it does not seem to be fully clear on which cortical level the convergence of the visual and proprioceptive input takes place. It has been proposed before that multisensory integration takes place at multiple sites within the nervous system [63], [64], [65].

It is conceivable that the different inputs are converging before being referenced to frontal mirror areas or M1 as proposed by the results of Barraclough et al. [66]. They identified neurons in the superior temporal sulcus (STS) which potentially form multisensory representations of perceived actions. A study on nonhuman primates also found neurons that exhibit signs of multisensory interactions with super-additive or sub-additive response summation located in the superior temporal region and spatially clustered [67]. In addition, several studies using functional magnetic resonance imaging (fMRI) in humans and nonhumans identified regions in the premotor cortex [31], [68] or posterior parietal cortex [69] with clusters of neurons exhibiting also polysensory responses which might be involved in the integration of multimodal information about action. Furthermore, in animal studies, parts of the midbrain, i.e. deeper laminae of the superior colliculus, have been found to contain multisensory neurons of all possible combinations [48], [70]. Thus, the superior colliculus was described as a primary site for the integration of converging inputs from multiple sources but not as a reflection of the multisensory integration process [71]. It is itself in so far not automatically capable of integrating these multiple sensory inputs but obviously relies on the projection from the association cortex for that purpose [71].

It was proposed that response enhancement is a general property of multisensory integration [72]. The fact that we found an additive multisensory enhancements speaks in favour of true multisensory integration. However, the interpretation of cumulative multisensory responses and the role of movement related signals is still discussed [73]. It has been suggested that superadditive responses should exceed additive responses reflecting the activity of neighbouring neurons of unisensory character as multisensory response interactions were found on a cellular level to be more a multiplicative than a summative change in activity [74], [75].

The meaning of cross-modal sensorimotor integration for motor control is obvious. However, the exact function of kinaesthetic information processing is currently still not completely understood. A mirror-like functioning of the kinaesthetic system might indeed be conceivable if one follows the extended mirror system concept of Rizzolatti et al. [76]. For example, kinaesthetic training is common in sports such as dance and tennis, when the trainer provides the “feel” of the correct posture and movement by manually guiding the leg or hand.

One part of the kinaesthesis is the contribution of sensors as Golgi tendon organ, cutaneous stretch receptors and the muscle spindle receptor afferents to the sensory cortex. In the present study, we are interested specifically in the importance of different information in motor control. Certainly one can note that the kinaesthetic information is of considerable significance on movement execution and motor control. A disturbance of kinaesthesia has an obvious effect on the execution quality and precision of movement. Both need an undisturbed awareness of the peripersonal space which is the space within reach. As more recently proposed there are different spatial representations as to the result of motor interaction with the environment depending on the different effector properties [77]. It can be assumed that the emergence of these representations relies on multimodal sensory input. In this context, it has been found that the core part of the cortex involved in the encoding process of the peripersonal space is premotor area F4 [78]. Further, the ablation of the ventral part of the premotor cortex in a study with monkeys resulted in a so-called peripersonal neglect [79] which in turn hints at disturbed integrative processing of the different modalities including kinaesthetic information.

The cross-modal interaction between extero- and proprio-feedback is used to optimize information for motor control. In our experiment there is some evidence, that kinaesthetic information about limb movement, provided by a motor-driven wrist flexion-extension-machine, has a greater influence on the task-specific motor circuits than the visual information about the same movement.

## 4. Conclusion

In conclusion, multimodal sensory integration is fundamentally important for motor control. The present data complement the current view of the neural basis of the integration process within the cerebral motor circuits. In our experiment, we investigated the link between two different input modalities, i.e. visual and kinaesthetic information, and the influence of its combination on the motor outcome. The study provides evidence that the motor system excitability is significantly enhanced in subjects that perceive simultaneous information of both modalities about the movement. As the excitatory corticospinal drive was less enhanced when only a single information modality was provided this hints at an information-specific modulation of corticospinal excitability. The results suggest the existence of “kinaesthetic motoneurons” that respond to kinaesthetic information induced even solely by passive movements. It furthermore supports the assumption that the process of merging multimodal information including kinaesthetic input is a general characteristic of the motor system.

## Author Contributions

Conceptualization, Volker Zschorlich and Frank Behrendt; Methodology, Volker Zschorlich, Frank Behrendt and Mark de Lussanet; Writing – review & editing, Volker Zschorlich, Frank Behrendt and Mark de Lussanet. All authors contributed to the analysis and interpretation of the data. All of the authors approved the final version of the manuscript.

## Institutional Review Board Statement

The study was conducted according to the guidelines of the Declaration of Helsinki, and approved by the Institutional Ethics Committee of the University Medicine Rostock (No. A 2016-0138).

## Informed Consent Statement

Informed consent was obtained from all subjects involved in the study.

## Data Availability Statement

Please refer to suggested Data Availability Statements in section “MDPI Research Data Policies” at https://www.mdpi.com/ethics.

## Acknowledgments

The authors are indebted to Dipl. Eng. Norbert Wolff and Dipl. Eng. Andreas Mattke for their technical support and building the wrist machine.

